# A whole organism screening platform identifies gut microbiome microproteins that modulate host metabolism

**DOI:** 10.64898/2026.02.11.705328

**Authors:** Nina E. Short, Eleanor C. Warren, Imani E. Porter, Yishay Pinto, Javier Rodriguez, Karen Sarkisyan, Ami S. Bhatt, Andre E. X. Brown, David T. Riglar

## Abstract

Advances in metagenomic sequencing over the past two decades have identified vast numbers of previously uncharacterised small open reading frames that may encode microproteins (<50aa). Although computational tools have accelerated gene sequence prediction from metagenomic data, the function of most annotated proteins remains unknown and untested, especially in the context of host-microbiome interactions. Here, we present a scalable phenotypic screening pipeline to identify gut microbiome-derived proteins that modulate host function. Using the nematode worm *Caenorhabditis elegans* as a whole animal model that is amenable to systematic screening approaches, our pipeline integrates high-throughput cloning, expression and delivery to worms via feeding, followed by behavioural phenomics screening. We apply this approach to a pilot library of 126 uncharacterised microproteins (< 50 aa) from healthy human gut metagenomes, identifying a set of high-interest targets with potential activity and ultimately validating a microprotein that modulates host fatty acid metabolism when expressed. With protein-based therapies increasingly recognised as a promising alternative to traditional small molecules, this work highlights the potential of a target-agnostic approach for the systematic screening and discovery of novel bioactive proteins.

## Introduction

Microbial microproteins, encoded by small open reading frames (smORFs) up to 150 nucleotides in length ^1,2^ are critical for microbial function ^3-8^. In recent years, improved tools for *in silico* and evidence-based smORF annotation ^9^ have led to systematic identification of putative smORFs ^10-17^. Owing to computational and experimental challenges arising from their small size, however, functional annotation of the encoded proteins remains sparse. The microproteins that have already been functionally characterised highlight diverse roles in microbial function, ecology and host interaction ^2^. Cytoplasmic or membrane-bound microproteins tend to be implicated in roles relating to cellular metabolism such as sporulation ^18,19^, cell division ^20^ and stress sensing and response ^21-24^. Secreted microproteins have been found to act as autoinducers, enabling inter-bacterial communication via quorum sensing ^25,26^, and mediate inter-microbial competition ^27-29^. Microprotein modes-of-action include recruiting larger proteins to the cell membrane, regulating protein stability, regulating protein complex formation and stability, and regulating membrane protein, soluble protein, and enzymatic activity ^30^.

To our knowledge, no bacterial microproteins have yet been directly implicated in host-microbe interactions. However, microbes are known modulate their host via several different mechanisms, with the host-microbe interface in the gut microenvironment being a particularly rich and widely researched site of bidirectional signalling. For example, various gut-resident bacterial species can produce physiologically relevant levels of the neurotransmitter gamma amino butyric acid (GABA) ^31^, with potential implications in host pathology ^32^. Proteins, and peptides derived from larger microbial genes, are also known to mediate microbe-host interactions. Heat-stable enterotoxins (ST) produced by enterotoxigenic *E. coli* (ETEC) mimic the structure and function of the human peptide hormone guanylin, activating the GC-C receptor on intestinal epithelial cells and inducing diarrhoea ^33^. Neurotoxins, including botulinum and tetanus toxins are also examples of bacterially-derived proteins with direct modulation of host neural pathways ^34^. Given the numerous cases of microbial-derived products impacting host physiology, the prevalence of bacterial molecular mimicry, and the importance of peptide signalling in the mammalian host, microproteins are thus strong candidates for mediating similar host modulation by microbes. However, novel systematic functional screens and targeted assays are needed to accelerate bioactive microprotein discovery efforts.

An ideal platform to identify bacterial microproteins capable of mediating microbiota-host interactions requires i) testing in whole animal context to reflect the variety of potential pathways for interaction, along with ii) high-throughput analysis and iii) rapid and affordable protein production capacities to reflect the magnitude of unknown smORFs in metagenomic datasets. To address this, we started with a dataset of approximately 450,000 putative smORFs derived from healthy human metagenomic data from the Human Microbiome Project (HMP) ^10^. To restrict this list to the strongest candidates for host impact in the gut microbiome, we selected only candidates expressed in gut-derived metagenomes and used computational tools to predict secretion. From this shortlist, we cloned a representative sample for heterologous expression in *E. coli*, yielding a library of 126 unique smORFs. To identify hits for more extensive follow up, we analysed these for impact on the nematode worm *C. elegans -* a well-established model system for diverse human functions and diseases that has emerged as a powerful tool for assessing small molecule bioactivity in varied drug discovery pipelines ^35,36^. We take advantage of C. elegans’ natural bacterivory; as the worms ingest and mechanically grind bacterial cells, the microproteins are released, analogous to the way in which feeding RNAi relies on bacterial ingestion to release dsRNA ^37^ Delivery by feeding facilitates microprotein exposure via cloned libraries without requiring engineered secretion from *E. coli*, a non-native host that may not export these microproteins efficiently, if at all. Neuropeptide delivery to loss of function *C*.*elegans* via bacterial feeding has previously been used in a low-throughput neuropeptide functional characterisation pipeline via a ‘rescue-by-feeding’ approach ^38^.

Our analysis builds on an existing *C. elegans* phenotypic screening platform, which combines high-resolution megapixel cameras ^39^ and automated worm tracking software ^40^ to acquire quantitative behavioural data. The approach has shown success in drug repurposing screens ^41^, modelling neurodegenerative diseases ^42^, and predicting the mode of action of insecticides and anthelmintics ^42^.

Applying this pipeline to functionally screen our pilot microprotein expression library, we identify several promising host-modulatory microproteins. Our platform therefore enables the systematic discovery of bioactive, smORF-encoded microproteins.

## Results

### Building a bacterial expression library of microbiome-derived bacterial microproteins

To initiate a systematic screen for bioactive microbiome-derived microproteins, we curated a subset of smORFs from the human gut microbiome, which we enriched for sequences with a greater likelihood of interacting with the gut microenvironment based on the criteria of being (1) gut-derived and (2) predicted to be transmembrane or extracellularly localised in their native host (Figure 1A). From a previously published dataset of ∼450,000 predicted microbial smORFs derived from healthy human metagenomes of the Human Microbiome Project (HMP) ^10^, we identified ∼150,000 sequences derived from gut metagenomic samples. Of these, 2,412 unique smORFs are predicted to contain Sec/SPI signal peptides by SignalP 5.0 software ^43^, and therefore have a higher potential of interacting with extracellular targets relative to intracellular microproteins. To test the concept, we selected a subset of these, chosen to represent diverse microprotein families ^10^ for cloning into *E. coli* to generate a pilot, IPTG-inducible bacterial expression library for downstream functional screening (Figure 1B; Supplementary Figure 1). The selected smORFs, codon optimised for expression in E. coli, were synthesised as an oligo pool and cloned into a pET9a expression vector. To facilitate tracking of expression while minimising the potential impact on short protein function, all proteins were C-terminally tagged with the 11 amino acid HiBiT tag. The resulting plasmid library was transformed into *E. coli* Lemo21, a strain optimised for expressing challenging proteins such as toxins or membrane proteins. Transformed isolates were arrayed into six 96-well microplates. Nanopore long-read sequencing of the arrayed library confirmed the presence of at least 126 unique smORFs (Supplementary Data 1)

**Figure 1.**
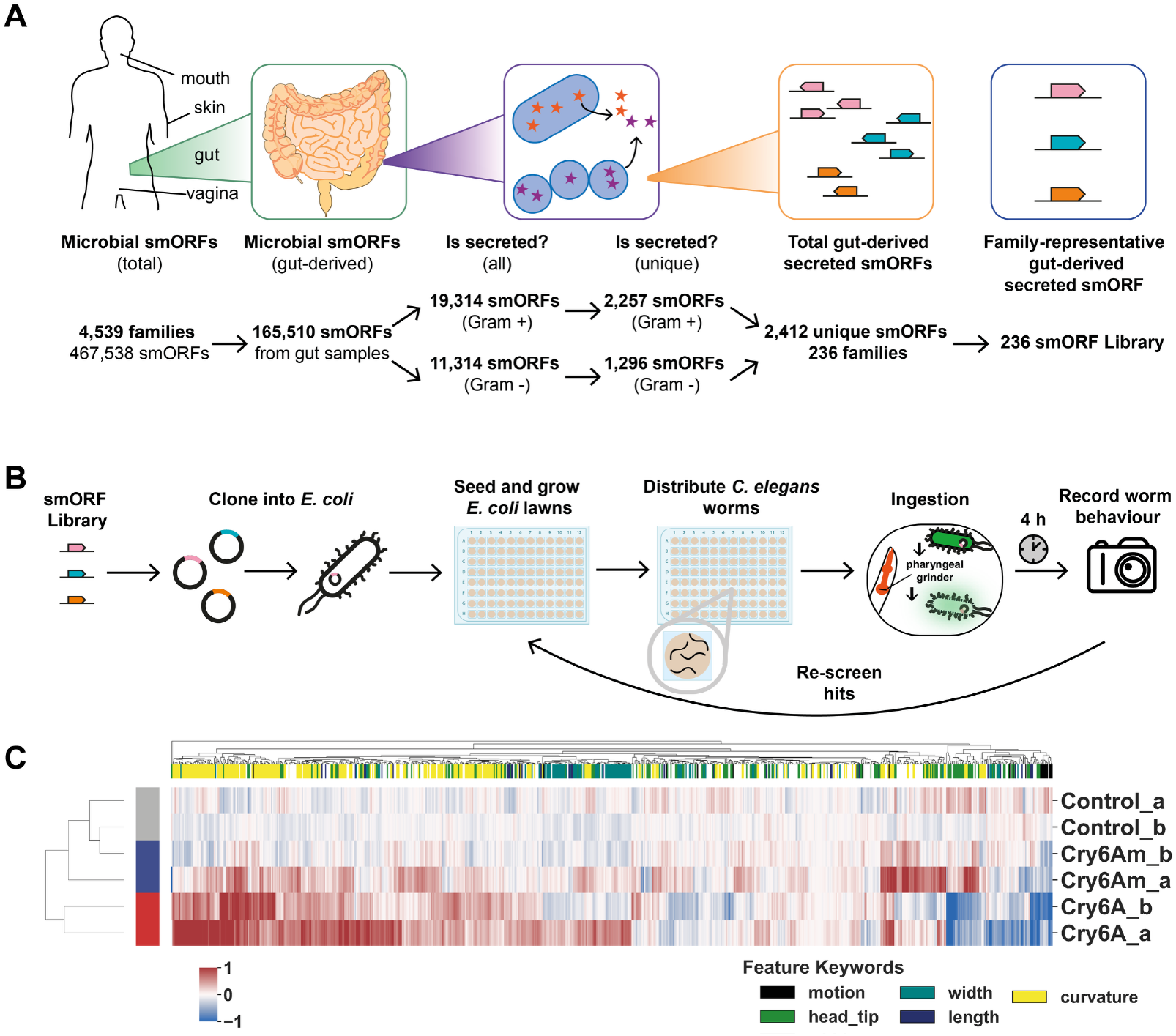
A pipeline for phenotypic screening of gut-derived bacterial microproteins in *C. elegans*. **A**. A diverse 236-member library of gut-derived putative secreted microproteins was shortlisted via bioinformatics. From an existing dataset of microbial smORFs ^10^, those derived from gut metagenomic samples and predicted to be secreted were selected. Removal of redundant sequences resulted in a list of 2,412 total proteins. Representatives from each microprotein family were chosen to maximise protein diversity **B**. The smORF library was synthesised as an oligo pool, cloned into a pET9a expression vector using pooled SapI-mediated Golden Gate Assembly, then transformed into *E. coli* Lemo21. Positive control strains expressing the nematicidal toxin Cry6A and a reduced-function mutation, Cry6Am, were also produced. The resulting cloned expression library was arrayed into 96-well microplates, which included strains expressing 126 unique microproteins. To screen, bacteria were grown as lawns and Day 1 adult *C. elegans* N2 worms were allowed to crawl freely across the lawn. After 4 hours of exposure, worm behaviour was recorded for ∼16 minutes and quantified using a semi-automated worm tracking and statistical analysis pipeline. **C**. Hierarchical clustering of worm behavioural fingerprints in response to control, Cry6A, or Cry6Am exposure. Number of replicate wells screened: Cry6A_a: 43; Cry6A_b: 159; Cry6Am_a: 49; Cry6Am_b: 104; Control_a: 47; Control_b: 264. Each column corresponds to an individual feature from the Tierpsy 48 feature set, and each row shows the average Z-normalised feature value for that feature across all replicate wells of given condition. Rows were clustered using average linkage hierarchical clustering with Euclidian distance computed across all features of the matrix. Columns are coloured based on whether the feature name contains one of the key words listed below the heatmap. Rows are coloured according to the experimental condition.

Two positive controls were cloned into the same expression vector and transformed into *E. coli* Lemo21: Cry6A, a nematicidal toxin derived from *Bacillus thuringiensis* that is known to impact *C. elegans* growth, locomotion, reproduction and survival ^44^; and Cry6Am, a mutant variant with a single amino acid substitution that reduces its toxicity to nematodes ^45^.

### Bacterial expression for delivery of host-modulatory proteins

To investigate the impact of this bacterial expression library in a whole animal context, we leveraged a high-throughput semi-automated phenotyping assay in the nematode worm *C. elegans*, which can capture worm behavioural responses to genetic perturbations and environmental stimuli (Figure 1B, Supplementary Figure 2) ^39,41^. A core set of morphological and behavioural features (e.g. morphology, posture, velocity) are extracted from videos of freely behaving worms and are then systematically expanded via a series of operations including localisation to body segments, calculating derivatives of time-series data, and subdivision according to worm motion state (Supplementary Figure 2) ^40^.

We delivered recombinant microprotein-expressing strains to *C. elegans* as bacterial lawns. Worms ingest and mechanically grind bacterial cells and are thus exposed to intracellular and secreted bacterial proteins (Figure 1B). As expected from previous studies, delivery of control Cry6A via feeding on agar plates impacted worm survival and motility while the Cry6Am mutant had reduced impact (Supplementary Figure 3) ^44,45^. Similarly, when worms were exposed to recombinantly expressed Cry6A and screened via high-throughput behavioural phenomics, they demonstrated considerable behavioural impacts compared with control conditions (631 and 3458 behavioural features significantly impacted in two independent screens). By comparison, exposure to Cry6Am-expressing bacteria impacted fewer features (32 and 431 significantly impacted features). Consistent with Cry6A’s expected role, impacted features suggested impaired locomotion, an increase in the amount and rate of worm bending/coiling behaviour, a reduction in worm length, and a reduction in exploratory head movement and reversal behaviour (Supplementary Figure 4).

Hierarchical clustering of behavioural ‘fingerprints’ across screens was able to clearly separate Cry6A exposed worms from those of control conditions or the mutant variant, Cry6Am (Figure 1C). Together, these results validate high throughput recombinant bacterial lawns as an effective delivery mechanism when screening proteins for bioactivity in *C. elegans*.

### Shortlisting of microproteins with potential behavioural impact

To shortlist microproteins from our expression library with behavioural impacts, we employed a sequential screening approach (Figure 1B). Two rounds of screening, initially of the full library (Screen 1) and subsequently of a subset of top responders (Screen 2), identified isolates significantly impacting at least one behavioural feature of *C. elegans* (Screen 1: 186 of 565 total and Screen 2: 60 of 186 total). Wells with poor bacterial growth were eliminated from the analyses, and the was no correlation seen between bacterial isolate growth rate and the number of significant features impacted by that isolate suggesting that hits were not driven by bacterial growth or lawn density factors (Supplementary Figure 5). Sequencing identified isolates expressing 41 unique microproteins within this initial list of hits (Supplementary Data 2).

We used a combination of factors to further prioritise hits for follow-up assays, including total number of features impacted, evidence of shared features impacted across screens, and independent identification of replicate isolates from the initial library. Based on these criteria, we selected 9 microproteins for further investigation (Figure 2A). To reduce the chance of false positives, we quantified the growth and expression profiles of all isolates identified expressing these 9 microproteins following 24 hours growth on solid media with or without 0.5mM IPTG to induce microprotein expression (Figure 2B-C). Comparison of isolate growth on agar identified several isolates with reduced growth in the presence of IPTG, as expected from the additional metabolic burden caused by heterologous protein expression (Figure 2C). Both isolates expressing microprotein 386515 and the sole isolate expressing microprotein 391283 showed no evidence of growth in the presence of IPTG and so were excluded from follow up screens. Microprotein expression levels were also quantified using a HiBiT blotting assay following overnight growth on blotting paper overlaid on LB agar, cell lysis and transfer to PVDF membrane. There was no evidence of expression for any isolates for microproteins 368440, 377726, 391283, or 156126 (Figure 2C). Selecting only isolates with evidence for both growth and expression led to a final shortlist of 4 high-priority lead smORFs (Figure 2D).

**Figure 2.**
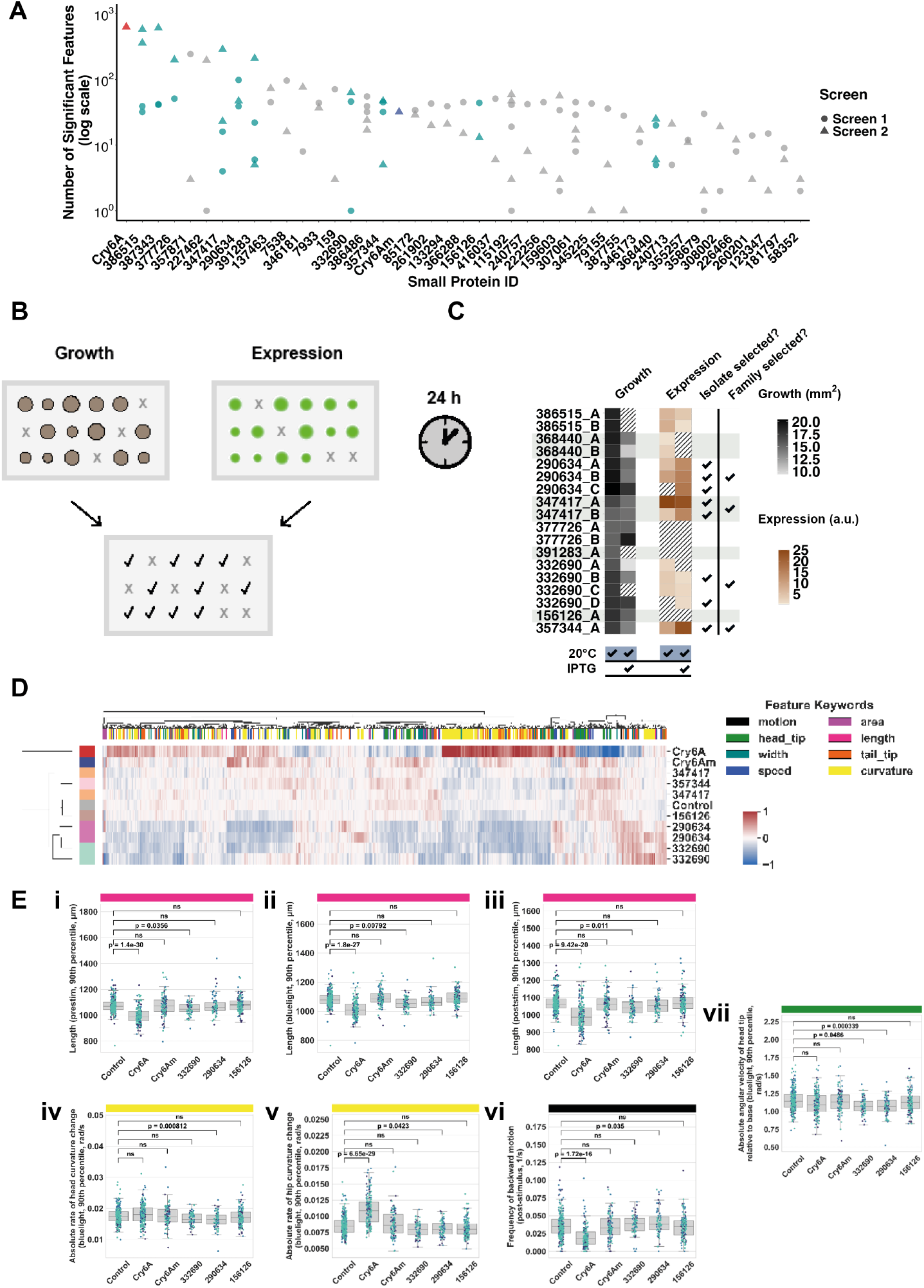
Shortlisting microprotein candidates with potential behavioural impacts. **A**. Number of significantly impacted features after exposure to a given recombinantly expressed microprotein. Shapes correspond to screen, and shortlisted strains of interest are highlighted in teal. **B**. Schematic of growth and expression assays. **C**. Results of the growth and expression assay. Heatmaps show (left) average colony size (mm^2^) and (right) average integrated density (arbitrary units) following a HiBiT blotting assay on lysed cells. Both assays were undertaken after 24 hours growth at room temperature on LB agar ± 0.5mM IPTG, with n=2 independent repeats. Hatched cells denote samples where growth/expression was not detected. **D**. Hierarchical clustering of worm behavioural fingerprints, as calculated from n=3 independent days of testing with number of n=x-y wells tested per isolate ranging from 32 to 138 (for microprotein-expressing strains) and 104 to 264 (for controls). Each column corresponds to an individual feature, and each row shows the average Z-normalised feature score for that feature across all replicate wells of given condition. Rows were clustered using average linkage hierarchical clustering with Euclidian distance computed across all features of the matrix. Columns are coloured based on whether the feature name contains one of the key words listed to the right of the heatmap. Rows are coloured according to the experimental condition. **E**. Boxplots showing distribution of scores for representative features across multiple assay conditions. Each point corresponds to a single well, and the score is averaged across all worms (approximately 3 worms per well) present. For each feature, scores from the control group were compared to those from each environmental condition using univariate two-sample t-tests. P-values were corrected for multiple comparisons using the Benjamini–Yekutieli procedure to control the false discovery rate. ns: not significant (p > 0.05). Colours indicate independent data taken across 3 days.

### Analysis of microproteins 332690 and 290634

To further investigate the impact of microproteins 332690 and 290634 independent of recombinant *E. coli* expression and HiBiT tags, we next measured the impact of chemically synthesized microproteins on *C. elegans* behaviour. Assay plates were seeded with lawns of the standard *E. coli* control strain, adding synthetic microprotein corresponding to the predicted mature forms of each protein at a final concentration of 10 µM, 1 µM or 0.1 µM prior to dispensing the worms. After 4 hours of exposure, we assessed *C. elegans* response to the microproteins using the standard analysis pipeline. Across all microproteins, Spearman correlations between synthetic and recombinant versions ranged from 0.275 to 0.438, indicating weak to moderate positive correlations, with (Supplementary Figure 6). microproteins 290634 and 332690 showing the strongest correlation coefficients (0.426 and 0.438, respectively). However, considerable unexplained variation remains that may result from variability in peptide solubility, concentrations delivered to the gut by either method, peptide maturity or folding differences, or various other differences between the recombinant and synthetic assays. Nevertheless, we measured 18 behavioural features that were impacted by both isolates of recombinantly expressed microprotein 332690 and at least one concentration of the synthetic microprotein (3.7% - 21% of total significant features of each isolate), and 4 features were shared between microprotein 290634 and one concentration of the corresponding synthetic microprotein (9.5-66.7% of total significant features of each isolate). In all cases, the directionality of the impact of the recombinant and synthetic microproteins relative to the control were the same (Supplementary Figure 7).

Whilst limited information is available about the two shortlisted smORFs (Figure 3A), both were predicted by AlphaFold2 ^46^ to encode alpha-helical proteins as full length proteins and unstructured or alpha-helical in predicted mature form (Figure 3B), although Alphafold’s limitations for such short proteins and propensity to overestimate alpha-helical tendencies in short proteins must be taken into account here ^7,47^.

**Figure 3.**
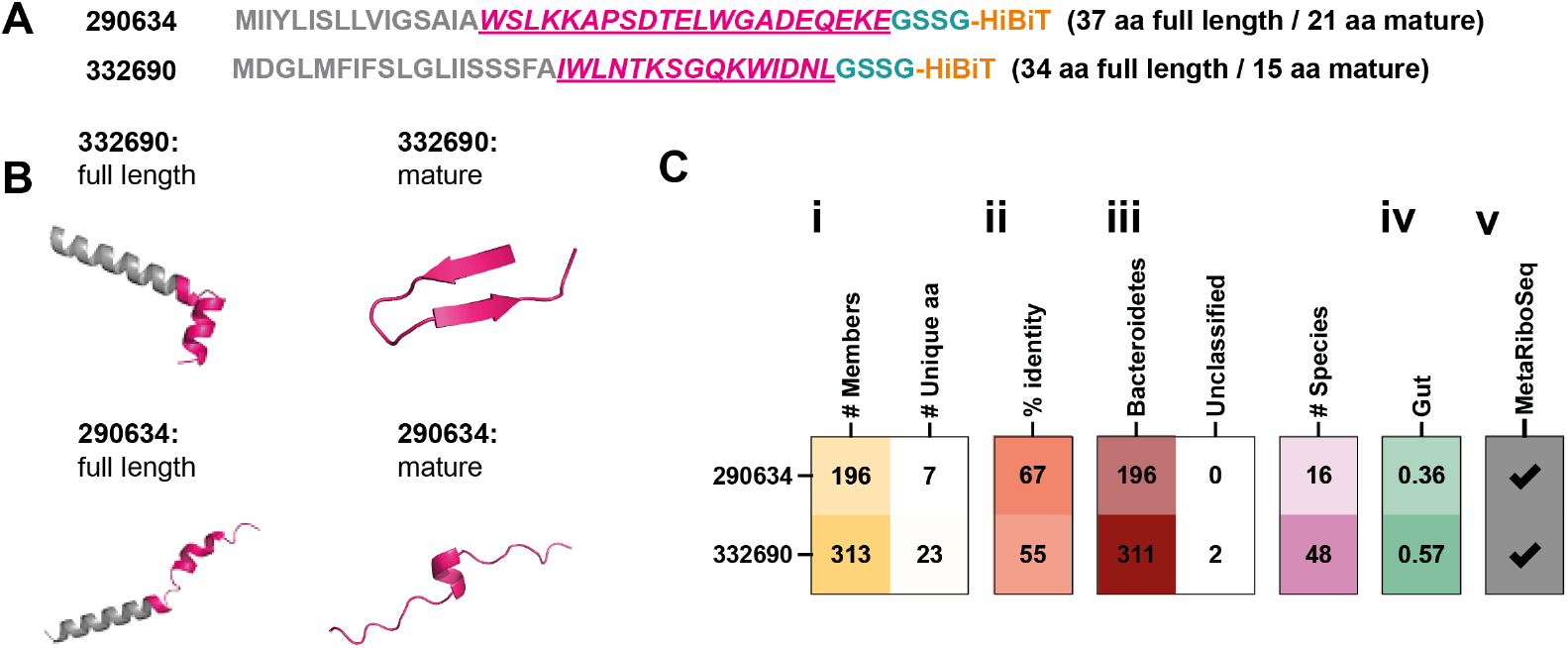
Bioinformatic attributes of shortlisted microprotein families 332690 and 290634. **A**. Sequences of microproteins 290634 and 332690 contained in our library, with predicted mature sequence highlighted. **B**. Alphafold2 predicted full length and mature sequences for each microprotein. Isoelectric points, full length / mature: 332690 (6.031 / 9.177), 290634 (4.038 / 4.038). C. General properties of the broader microprotein families 290634 and 332690 including i) number of other members, ii) full length sequence identity (including predicted signal peptide sequence) across unique family members, iii) Phyla of family members and number of species represented; iv) percentage of gut metagenomes with a representative from the family and v) prediction for active translation based on MetaRiboSeq analysis. Data are derived from ^10,48^.

Expanding the analysis to the level of the smORF families, as defined by CD-Hit clusters ^10^ (Figure 3C), demonstrated that members of both 332690 and 290634 families were almost exclusively found within Bacteroidota, with 332690 family members in 48 species and 290364 in 16. Members of the family to which microproteins 332690 and 290634 belong were detected in 57% and 36% of gut metagenomic samples, respectively. Members of both these families were also previously predicted to be translated in the gut using the ribosomal profiling tool MetaRibo-Seq ^48^. Taken together, these data suggest we have identified microproteins that are expressed in the gut within their native context. 332690 family members are also regularly found near horizontal-gene transfer (HGT) associated genes ^10^, and more recently homologues have been annotated by Cenote-Taker 2 viral discovery and annotation tool to map to bacteriophages of the class Caudoviricetes ^49^. It is thus likely the smORF is encoded by a prophage within the genomes of Bacteroidota.

### Recombinantly-expressed microprotein 332690 impacts *C. elegans* fatty acid metabolism

To further confirm and investigate microprotein 332690 and 290634 bioactivity, we assessed their impact on the *C. elegans* transcriptome (Figure 4A). We again included the microprotein 156126-expressing strain as a negative control. Following 4 hours of exposure to the different bacterial strains, we extracted and sequenced bulk *C. elegans* RNA. Growth on *E. coli* expressing microproteins 156126 and 290634 led to minimal differentially expressed genes (DEGs) and no functional enrichment (Figure 4B). By comparison, growth on *E. coli* expressing microprotein 332690 showed extensive changes to gene expression in the worms (Figure 4B). Across 2 assays in different worm strains and different facilities, we identified consistent modulation of 15 genes (Figure 4C, Table 1), with functional enrichment of genes related to fatty acid metabolism (Figure 4D), many of which are known to be expressed in the *C. elegans* intestine.

**Figure 4.**
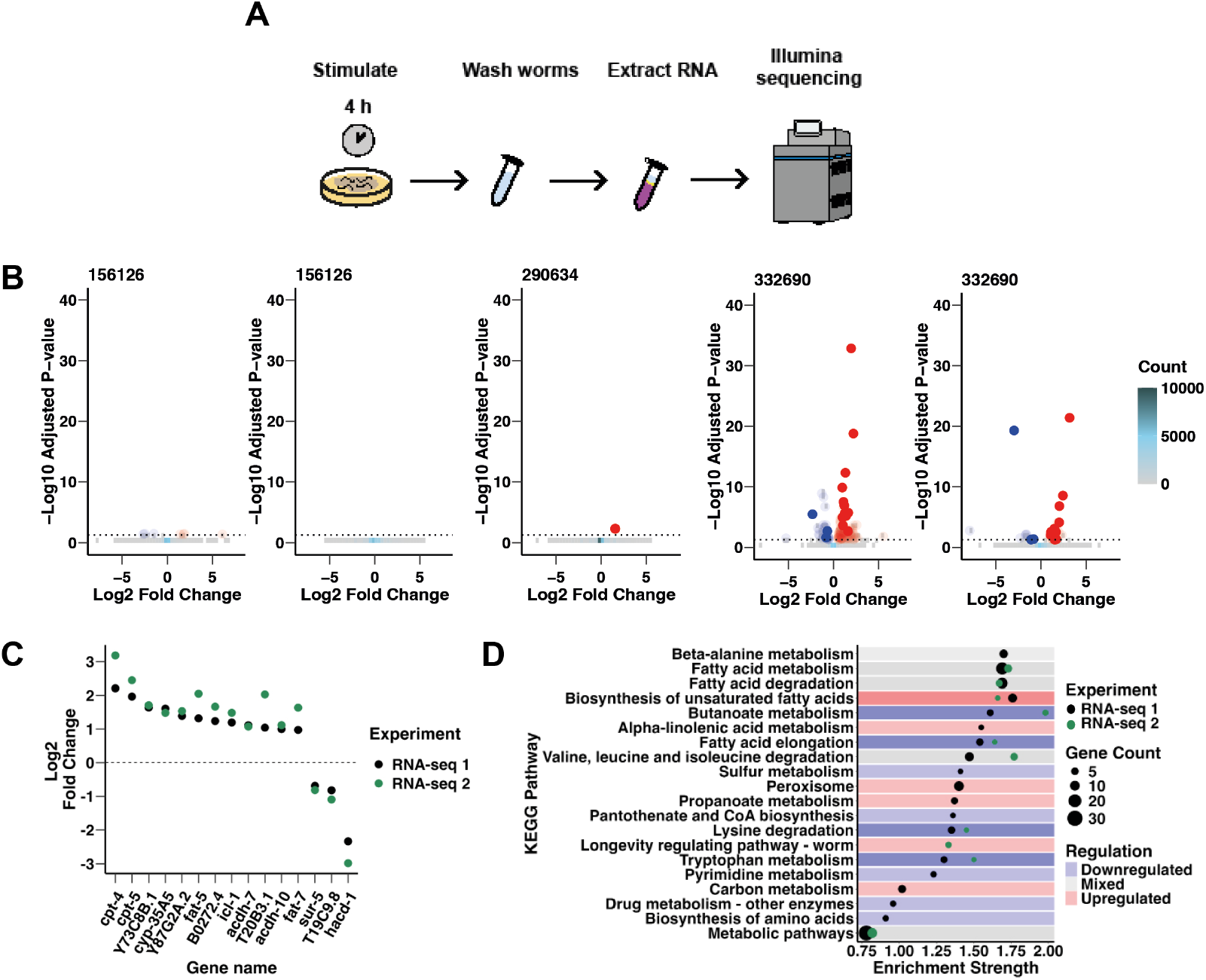
Recombinantly expressed microprotein 332690 impacts *C. elegans* fatty acid metabolism. **A**. Overview of experimental pipeline to assess recombinantly expressed microprotein impact on the *C. elegans* transcriptome. Worms were stimulated on a bacterial lawn for 4 hours at 20°C, after which they were washed off plates with M9, and RNA was extracted and sequenced by Illumina short-read sequencing. **B**. Genes significantly differentially expressed in response to recombinantly expressed microprotein exposure across two screens. RNA-seq 1: two plates per experimental condition, with ∼150-200 worms per late. RNA-seq 2: three plates per experimental condition, with ∼250-300 worms per plate. Genes were considered significantly differentially expressed if their p-adjusted value was < 0.05. **C**. Magnitude of differential gene expression in 15 genes significantly impacted in both assays with 332690 exposure. Points are coloured according to the assay of origin. **D**. KEGG pathways that are significantly enriched in the 106 and/or 33 genes that were differentially regulated by microprotein 332690 across both RNA-seq assays according to STRING analysis ^50^. All results shown are statistically significant with p-values corrected for multiple comparisons using the Benjamini-Hochberg procedure (FDR < 0.05). Points are coloured according to the source assay, and their size is proportional to the number of genes associated with a given KEGG pathway. Rows are coloured according to whether the genes annotated with each term were all upregulated in both assays (red), all downregulated in both assays (blue), or both up and down regulated in both assays and/or within the same assay (grey).

**Table 1:**
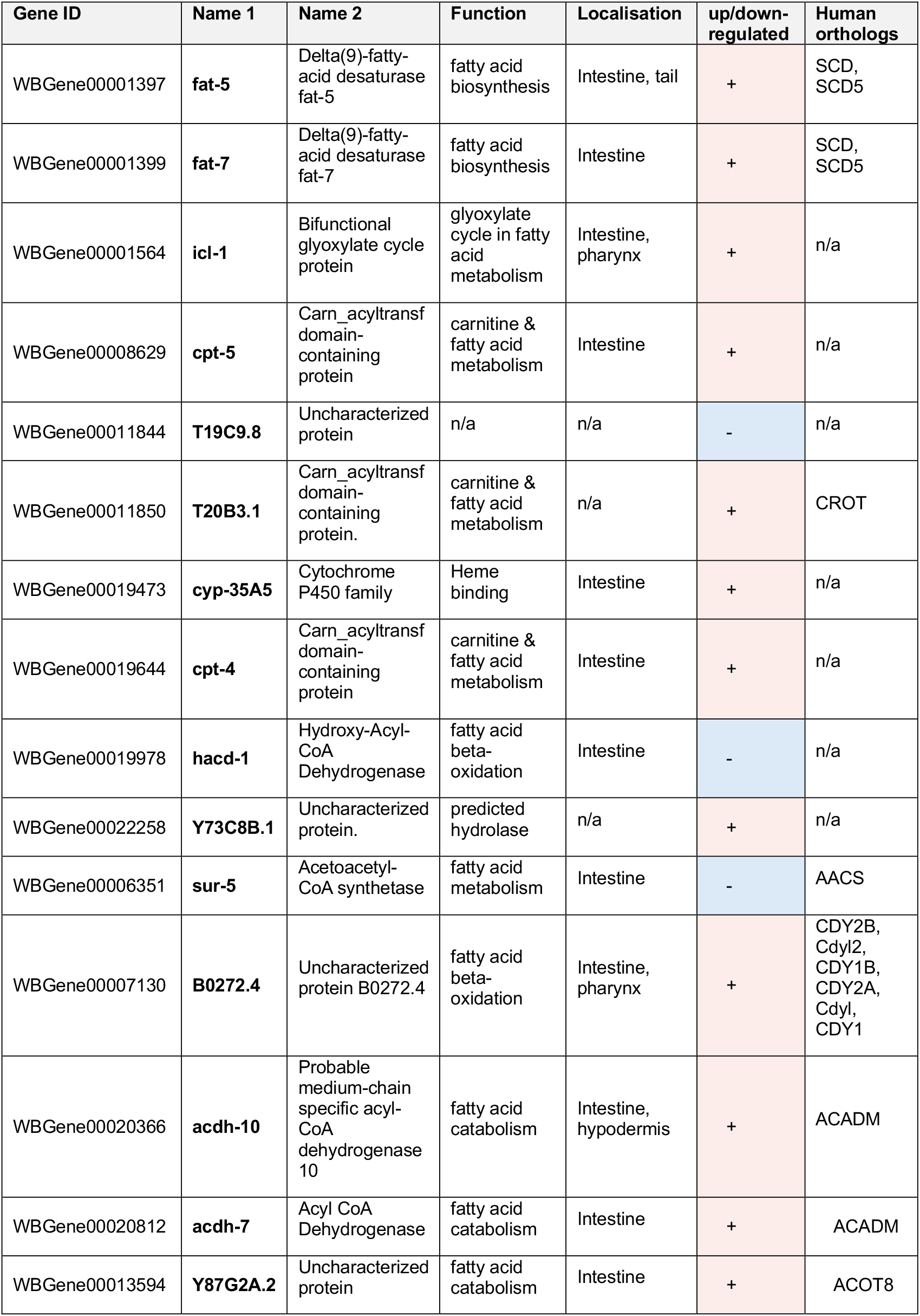
15 C. elegans genes consistently impacted by recombinantly expressed microprotein 332690.

| Gene ID | Name 1 | Name 2 | Function | Localisation | up/down-regulated | Human orthologs |
| --- | --- | --- | --- | --- | --- | --- |
| WBGene00001397 | **fat-5** | Delta(9)-fatty-acid desaturase fat-5 | fatty acid biosynthesis | Intestine, tail | + | SCD, SCD5 |
| WBGene00001399 | **fat-7** | Delta(9)-fatty-acid desaturase fat-7 | fatty acid biosynthesis | Intestine | + | SCD, SCD5 |
| WBGene00001564 | **icl-1** | Bifunctional glyoxylate cycle protein | glyoxylate cycle in fatty acid metabolism | Intestine, pharynx | + | n/a |
| WBGene00008629 | **cpt-5** | Carn_acyltransf domain-containing protein | carnitine & fatty acid metabolism | Intestine | + | n/a |
| WBGene00011844 | **T19C9.8** | Uncharacterized protein | n/a | n/a | - | n/a |
| WBGene00011850 | **T20B3.1** | Carn_acyltransf domain-containing protein. | carnitine & fatty acid metabolism | n/a | + | CROT |
| WBGene00019473 | **cyp-35A5** | Cytochrome P450 family | Heme binding | Intestine | + | n/a |
| WBGene00019644 | **cpt-4** | Carn_acyltransf domain-containing protein | carnitine & fatty acid metabolism | Intestine | + | n/a |
| WBGene00019978 | **hacd-1** | Hydroxy-Acyl-CoA Dehydrogenase | fatty acid beta-oxidation | Intestine | - | n/a |
| WBGene00022258 | **Y73C8B.1** | Uncharacterized protein. | predicted hydrolase | n/a | + | n/a |
| WBGene00006351 | **sur-5** | Acetoacetyl-CoA synthetase | fatty acid metabolism | Intestine | - | AACS |
| WBGene00007130 | **B0272.4** | Uncharacterized protein B0272.4 | fatty acid beta-oxidation | Intestine, pharynx | + | CDY2B, CdyI2, CDY1B, CDY2A, CdyI, CDY1 |
| WBGene00020366 | **acdh-10** | Probable medium-chain specific acyl-CoA dehydrogenase 10 | fatty acid catabolism | Intestine, hypodermis | + | ACADM |
| WBGene00020812 | **acdh-7** | Acyl CoA Dehydrogenase | fatty acid catabolism | Intestine | + | ACADM |
| WBGene00013594 | **Y87G2A.2** | Uncharacterized protein | fatty acid catabolism | Intestine | + | ACOT8 |

**Table 2.**
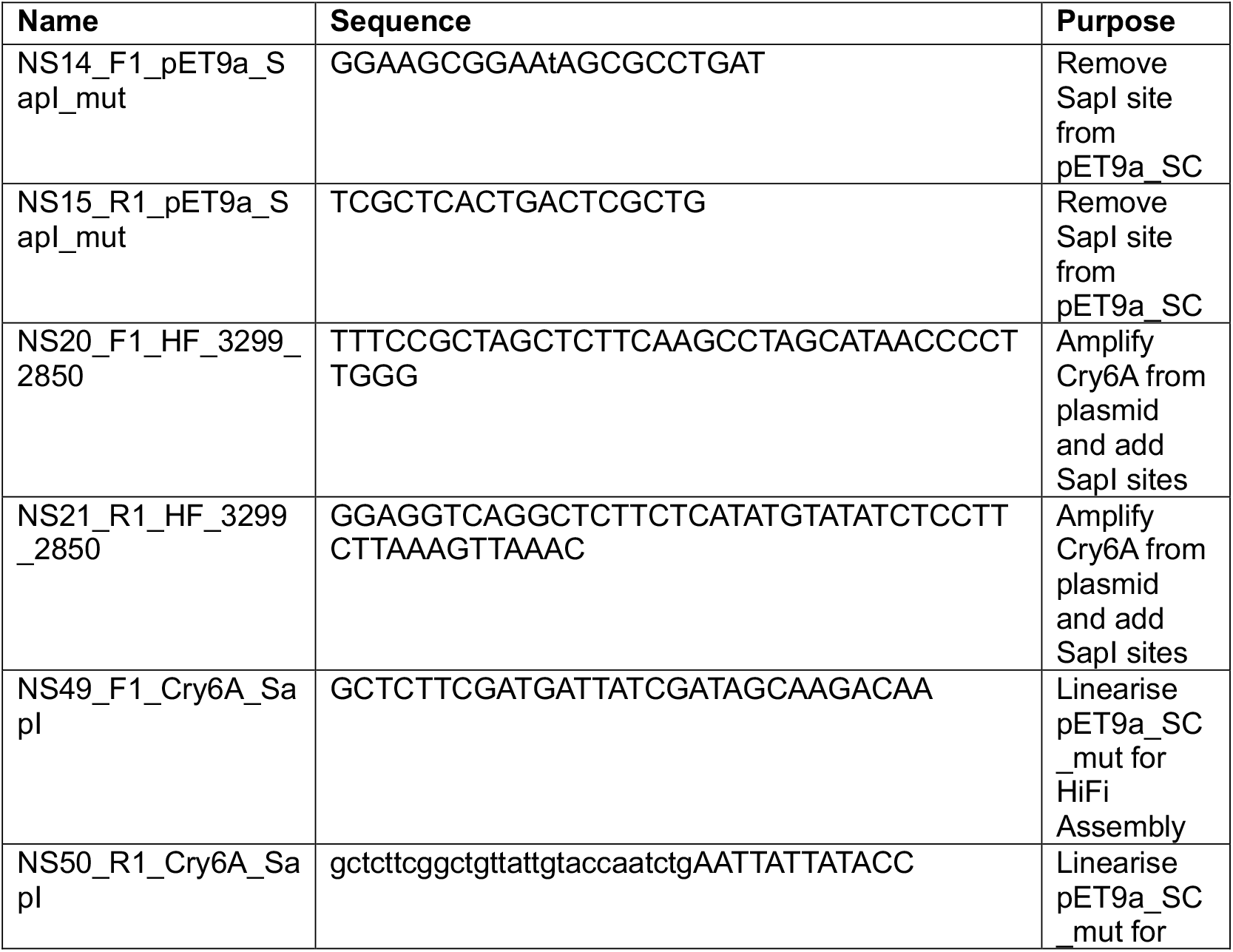
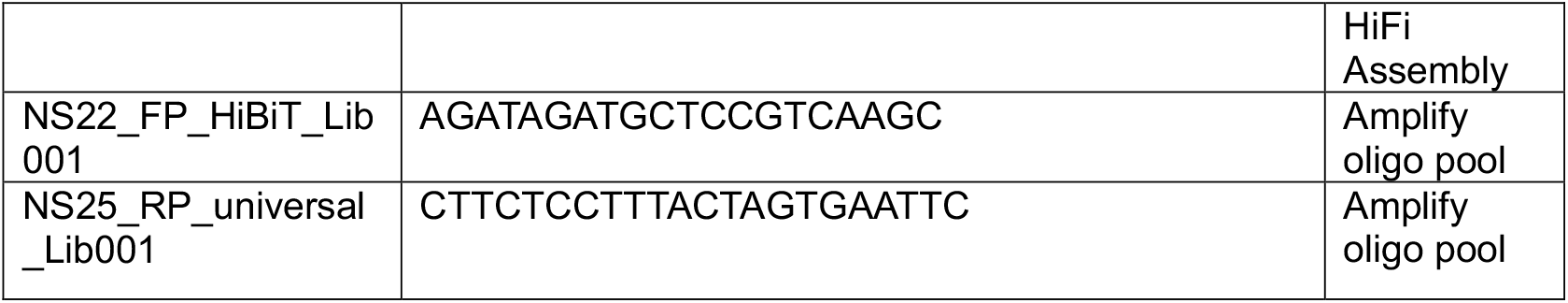
Primers used in this work.

**Table 3.**
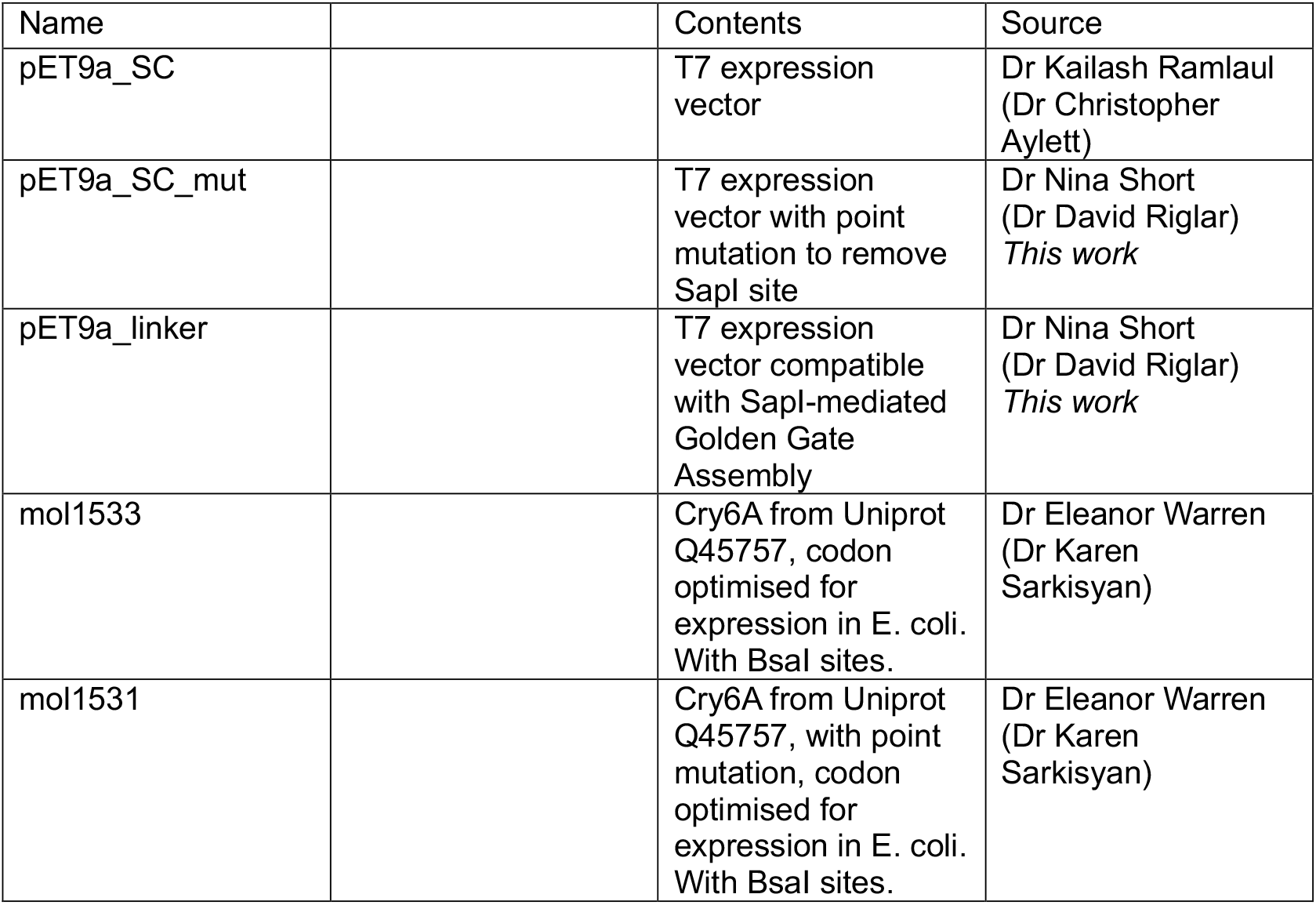
Plasmids used in this work.

| Name |  | Contents | Source |
| --- | --- | --- | --- |
| pET9a_SC |  | T7 expression vector | Dr Kailash Ramlaul (Dr Christopher Aylett) |
| pET9a_SC_mut |  | T7 expression vector with point mutation to remove SapI site | Dr Nina Short (Dr David Riglar) *This work* |
| pET9a_linker |  | T7 expression vector compatible with SapI-mediated Golden Gate Assembly | Dr Nina Short (Dr David Riglar) *This work* |
| mol1533 |  | Cry6A from Uniprot Q45757, codon optimised for expression in E. coli. With BsaI sites. | Dr Eleanor Warren (Dr Karen Sarkisyan) |
| mol1531 |  | Cry6A from Uniprot Q45757, with point mutation, codon optimised for expression in E. coli. With BsaI sites. | Dr Eleanor Warren (Dr Karen Sarkisyan) |

## Discussion

Microproteins are an underexplored class of bacterial product. Given the increasing evidence that they play important roles in bacterial physiology and ecology, characterising bacterial microprotein function is not only of basic biological importance but could also be harnessed for translational applications. Indeed, there is a wider growing interest in mining naturally derived proteins and peptides for therapeutic development, in contrast to traditional drug discovery efforts which tend to be centred around small molecules. Clinically relevant examples include cone snail neurotoxin-derived ziconotide, used to treat chronic pain ^51^ botulinum neurotoxin, used in treatments for diverse neurological disorders as well as cosmetic applications ^52^, and the glucagon-like peptide 1 receptor agonist semaglutide (Ozempic/Wegovy), used for type-2 diabetes treatment and weight loss ^53^. Despite the abundance of metagenomic data from which increasingly comprehensive and refined databases of smORFs are being predicted, there is a lack of systematic screening platforms for functional characterisation of the microproteins they encode in the context of host-microbe signalling.

To address this, we developed a pipeline to screen recombinant microprotein expression libraries in *C. elegans* for bioactivity. When designing this pipeline, scalability was of utmost importance. For this reason, we prioritised automation and pooled approaches where possible. Although this contributed to a reduced library coverage, with only 126 of 236 microproteins represented in our 6×96 well arrayed pilot library and didn’t allow for individual strain optimisation such as microprotein expression level standardisation, these factors were outweighed by the efficiency and scalability afforded by the pooled approach. The ability to screen multiple conditions in parallel (up to 480 conditions per 16-minute image run if using 96-well microplates) and carry out multiple runs per day means that this pipeline could easily be scaled by an order of magnitude to screen larger libraries. Automated colony picking and bacterial culturing would further simplify the process of arraying and preparing overnight liquid cultures of assay strains and the affordability of pooled oligonucleotide synthesis is similarly compatible with scale up. In future iterations, a larger library size or arrayed cloning methods could mitigate for inherent inefficiencies of pooled cloning. Performing expression and growth validation only on isolates of interest as demonstrated here (Figure 2C) reduces the bottleneck of these lower throughput assays.

*C. elegans* is an established model for small molecule drug screens ^54,55^ and proved to be well adapted to this screening pipeline. Firstly, whilst heterologous proteins are notoriously difficult to secrete, our setup bypasses the need for protein secretion or purification by delivering proteins via lawn feeding. This simplifies the assay process to a level feasible for high throughput testing. Secondly, *C. elegans* enables assaying the whole-organism phenotypic read-out, something not feasible in standard cell-based assays. Thirdly, *C. elegans*’ fully sequenced genome, invariant cell lineage, and fully mapped connectome also facilitate downstream mechanistic assays.

Our pipeline reliably identified the effects of the known nematicidal toxin Cry6A and its less toxic variant Cry6Am. Cry6A belongs to a pore-forming family of crystal proteins whose nematicidal effects are mediated by the formation of toxin pores in host cells. Cry6A in particular has also been shown to interact with host intestinal cells via the RBT-1 receptor ^56^, and contribute to host cell death via the activation of the aspartic protease-dependent host necrosis signalling pathway ^57^. In our screening, this impact was evident in terms of the number of features impacted and the magnitude of the impact. Impaired worm locomotion supports our hypothesis that recombinantly expressed proteins can be delivered to *C. elegans* via a bacterial lawn, that the proteins can be ingested by and act on *C. elegans*, and that this impact can be detected as a distinct behavioural phenotype. By comparison to the known doses of Cry6A that are lethal (18.5 µg/mL) and sub-lethal but leading to uncoordinated locomotion (5-10 µg/mL), we can also estimate the dosage equivalent from the 4 hour exposure to be in the order of 10 µg/mL ^44^.

In contrast to the impact of Cry6A, the impact of our lead microproteins was less pronounced and therefore did not allow for a clear behavioural profile to be established. This is unsurprising given the provenance of the microproteins from healthy human microbiomes, and thus with phenotypes both potentially small and beneficial rather than detrimental. Nevertheless, the consistent impact on *C. elegans* behaviour across three replicate screens, each involving three independent days of screening representing tracking of >700 animals, and with separately cloned isolates independently identified from the initial library, our approach highlights two consistent hits: microproteins 332690 and 290634. The ability of our screening pipeline to shortlist strains with bioactive potential is further demonstrated by the measured impact of recombinant microprotein 332690 on the *C. elegans* transcriptome, which has been validated across two worm strains in different labs.

Genes impacted by growth on 332690-expressing bacteria were associated with various pathways, including fatty acid metabolism. It is notable that several of these genes are controlled by Nuclear Hormone Receptor 49 (NHR-49). NHR-49 is an important regulator of *C. elegans* metabolism and lifespan which is activated during various stresses including starvation, oxidative stress, infection and hypoxia. Genes associated with its role in regulating mitochondrial β-oxidation (*cpt-5, ech-1*.*1*), fatty acid desaturation (*fat-5, fat-7)*, non-lipid metabolism, starvation and innate immune response (*icl-1)* and longevity *(lipl-1)* were all significantly modulated in 332690-fed worms. Interestingly, the directionality of regulation for several genes (cpt-4, fat-7, hacd-1) is suggestive of a satiety response given that they are opposite to those seen during starvation ^58^ and ivermectin exposure ^59^.

Future characterisation will be required to clarify the mechanism of action of microprotein 332690. This highlights one of the challenges of screening in whole animal models with various potential pathways for activity. Indeed, microproteins could be acting directly on worms or indirectly via an impact on *E. coli*. In this context, the mechanism for biological impact of microprotein 332690 expression is of additional biological interest given its annotation to a bacteriophage genome, raising the possibility for a virus to be manipulating bacterial and animal hosts in turn.

With the development and validation of these methods it will now be possible to screen a wider array of small and larger hypothetical proteins for impacts on gut function and behaviour.

## Materials and methods

### Bacterial strains

The destination vector for bacterial expression assays in this work was a version of the T7 expression vector pET9a_SC, modified to be compatible with SapI-mediated Golden Gate Assembly. To modify the original pET9a_SC vector, an existing SapI site was removed by site-directed metagenesis. The pET9a_SC vector was amplified by Q5 PCR (New England Biolabs) using primers NS14 and NS15 to introduce the modified base, after which 1 µL of purified PCR product (Monarch PCR and DNA Cleanup Kit, New England Biolabs) was incubated for 1 hour at room temperature with 1 µL each of T4 DNA ligase, T4 DNA ligase buffer, polynucleotide kinase and DpnI (Thermo Fisher Scientific) to ligate pET9a_SC_mut and digest any remaining template DNA. After linearising the pET9a_SC_mut using primers NS20 and NS21 to introduce necessary overhangs, a linker sequence containing two SapI sites was introduced by HiFi Assembly (New England Biolabs), forming pET9a_linker.

The sequences for *Bacillus thuringiensis* toxin Cry6A and Cry6Am were obtained from plasmids mol1533 and mol1531. The Cry6A sequence in mol1533 was originally derived from UniProt (Q45757) and had been codon optimised for expression in *E. coli*. The Cry6Am sequence in mol1531 was identical to the Cry6A sequence, with the exception that the leucine (L) at position 259 was mutated to aspartic acid (D), reducing the protein’s toxic effects. The sequences encoding Cry6A and Cry6Am were amplified from plasmids mol1533 and mol1531 by OneTaq PCR (New England Biolabs) using primers NS14 and NS15 and cloned into pET9a_linker by SapI-mediated Golden Gate Assembly to form pET9a_Cry6A and pET9a_Cry6Am. These were transformed into E. coli Lemo21 (New England Biolabs).

### C. elegans strains

C. elegans N2 worms were maintained on nematode growth medium (NGM) agar plates seeded with E. coli Lemo21 (New England Biolabs) at 20°C.

### Survival and locomotion assay

Overnight cultures of E. coli (control), E. coli (Cry6A) and E. coli (Cry6Am) in LB were back-diluted with LB to OD600 = 1, and 0.5 mL of back-diluted culture was seeded in duplicate onto 60 LB NGM agar plates and left to dry overnight at room temperature. 10 x Day 1 adult worms were dispensed onto each plate, incubated at 20°C, and survival and locomotion were scored visually following 0, 24, and 48 hours incubation. Scoring was performed by recording the percentage of surviving and motile (i.e. performing forward and/or reverse locomotion) worms.

### Standard phenotyping assay

All phenotyping assays were carried out in 96 square-well microplates (Whatman). 200 µL sterile media (LB NGM agar + 0.5 mM IPTG) was dispensed into each well using a ViaFILL liquid handler (INTEGRA Bioscience Ltd), left to solidify at room temperature, and stored at 4°C until use. One day prior to imaging, plates were dried in a LEEC BC2 drying cabinet (LEEC Ltd) until they lost 3-5% of their weight. Immediately after drying, plates were seeded with 10 µL of the relevant bacterial overnight culture and left to dry overnight at room temperature. For experiment with synthetic peptides, 10 µL of resuspended synthetic microprotein was dispensed onto seeded plates ∼1-2 hours before the addition of worms for a final concentration (including agar) of 10 µM, 1 µM or 0.1 µM. Plates were sealed with parafilm to allow microprotein resuspension to diffuse throughout the agar. Synthetic microprotein sequences and the relevant solvent are shown in Table 4.

**Table 4.**
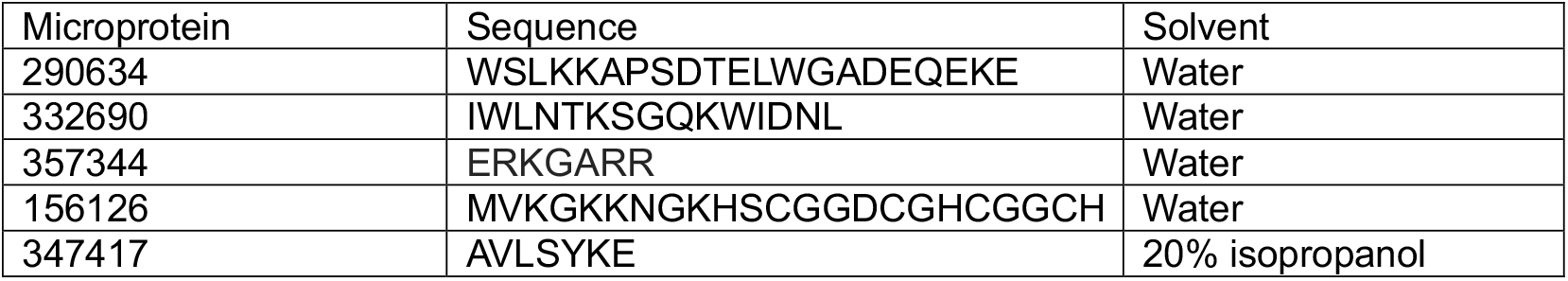
Synthetic microprotein sequences & solvents.

Day 1 adult worms were obtained by bleach-synchronisation, for which a detailed protocol can be found here: https://doi.org/10.17504/protocols.io.2bzgap6. On the day of imaging, Day 1 adult worms were dispensed onto imaging plates using either a COPAS 500 Flow Pilot wormsorter or a ViaFILL liquid handler (INTEGRA Bioscience Ltd). After drying in a sterile cabinet for 1-2 hours, plates were incubated at 20°C, then transferred to the imaging rooms 30 minutes prior to imaging to acclimate to the environmental conditions. Plates were imaged exactly 4 hours after the midpoint of the worm-dispensing step.

Plates were imaged using the Hydra Imaging Rig ^39^. The rig consists of 5 towers, each of which can image a single microplate via 6 x 12-megapixel cameras. A single imaging run can therefore image up to 5 microplates at a time. Each imaging run lasts 16 minutes and is subdivided into three phases: the pre-stimulus phase (5 minutes), bluelight phase (6 minutes, during which there are three bluelight pulses at 60, 160 and 260 seconds), and post-stimulus phase (5 minutes). Custom Python scripts with Loopbio’s API for Motif software were used to control automated plate recordings and photostimulation events.

The resulting raw video files were processed using Tierpsy Tracker software ^40^, which compresses videos, identifies and skeletonises individual worms, and identifies worm trajectories. For every feature analysed, a feature score is assigned to each worm in a single well, and then the average score across all worms in a well is taken. The resulting output is a feature matrix of average feature score per well where each row corresponds to a single well, and each column corresponds to a single feature.

Each raw video file was also annotated for quality control purposes. Wells were manually assessed, and any wells with issues such as fungal contamination, condensation, uneven agar, or anything else that could interfere with worm behaviour and/or tracking were flagged as ‘bad wells’.

When processing tracking data, several quality control steps were put in place to filter out poor or problematic data. Any wells annotated as ‘bad’ were dropped, as well as any wells with fewer than 100 tracked worm skeletons. Any wells with poor tracking (where > 40% of all features contained NaN or infinite values), or features with poor tracking (where > 20% wells contained NaN or infinite values) were also dropped from the annotated feature matrix.

In the event that a camera recording was out of focus, any associated recordings were also dropped from the final dataset. Cameras were determined to be out of focus if their median score for a representative width feature was 2 standard deviations or more above or below the median score for that feature across all cameras in the screen (Supplementary Figure 8A).

Results were also examined for day-to-day variation across screening days, by examining the distribution of feature scores for a given feature-set across all 3 days of screening (Supplementary Figure 8B). No day-to-day variation was found in any of the screens described in this work.

### Statistical analyses of standard phenotyping assays

Statistical analyses were carried using custom Python and R scripts. Univariate statistical tests were used to identify features that were significantly different in one experimental condition relative to the control condition. A false discovery rate (FDR) correction was applied using the Benjamini-Yekutieli method to minimise false positives resulting from multiple testing.

### Toxin phenotyping assay

Toxin phenotyping assays were based on the standard phenotyping assay described above. Two screens were carried out, each across three separate imaging days.

### smORF library bioinformatic pipeline

Custom Bash and Python scripts were used to subset the smORF dataset from Sberro and colleagues ^10^. SignalP 5.0 ^43^ was used to predict microprotein secretion status.

### smORF library cloning

smORF sequences were ordered as an oligo pool from Twist Bioscience. The oligo pool was resuspended in 10 mM sterile Tris buffer (pH 8) to 20 ng/µL. The pool was amplified by Q5 PCR using primers NS22 and NS25 with 14 cycles to minimise amplification bias. The purified PCR products (Monarch PCR and DNA Cleanup Kit, New England Biolabs) were cloned into pET9a_linker using SapI-mediated Golden Gate Assembly (New England Biolabs), and the resulting plasmids were transformed into E. coli Lemo21 and plated onto chloramphenicol/kanamycin selection plates. 565 colonies were arrayed into microplates to form Library 1. Library coverage was quantified by Nanopore sequencing. Library 1 was inoculated onto chloramphenicol/kanamycin selection plates, incubated overnight at 37°C, and the resulting colonies were washed off the selection plates and into a falcon tube using ice-cold LB, and miniprepped to retrieve pooled Library 1 plasmids (Monarch Plasmid Miniprep Kit, New England Biolabs). Pooled plasmids were linearised via EcoRI-mediated restriction digestion (New England Biolabs) and prepared for sequencing using a standard Ligation Sequencing Kit (SQK-LSK109, Oxford Nanopore Technologies). Nanopore sequencing was carried out on MinION flow cell, and Guppy was used for superaccurate basecalling of the resulting reads. Basecalled reads were then imported into Geneious Prime, trimmed using BBDuk, (trimming both ends, minimum quality Q7, discarding reads < 10 bp) and aligned to references using Minimap2.

### Growth and expression testing

To measure bacterial growth, 2μL overnight bacterial cultures were spotted onto LB agar plates containing 0 mM or 0.5 mM IPTG using a floating pin replicator (V& P Scientific). Plates were incubated at room temperature for 24 hours. Average lawn size (in mm^2^) was measured using a pipeline involving plate imaging using a Reshape Biotech plate imager and automated “small colony” size analysis using Reshape Discovery software.

Microprotein expression levels were measured using a spot assay. Overnight bacterial cultures were plated onto Whatman paper placed on the surface of LB agar plates containing 0 mM or 0.5 mM IPTG, and plates were incubated at room temperature for 24 hours. After lysing cells on the paper using 5% SDS, proteins were transferred from the paper to a PVDF membrane using an iBlot dry-transfer system. A HiBiT blotting assay was then carried out to quantify levels of HiBiT-tagged microprotein expression. Expression levels were quantified as the integrated density of the luminescent signal detected on membrane images.

### Microprotein phenotyping assay

The microprotein phenotyping assays were based on the standard phenotyping assay described above. In total, 3 assays were carried out, each taking place across 3 days. In Screen 1, Library 1 strains were screened. In Screen 2, all (i.e. strains impacting 1 or more features) from Screen 1 were screened. In Screen 3, a subset of hits from Screen 2 were screened. This sequential screening approach was used to narrow down the pool of microprotein candidates of interest.

### C. elegans transcriptomic assays

Two assays were conducted to examine the impact of recombinant microprotein expression on the C. elegans transcriptome. Both assays broadly followed the same protocol, with slight differences: RNA-seq 1 examined the impact of microproteins 332690 and 156126 on C. elegans, and featured 2 replicate plates per condition, whilst RNA-seq 2 examined the impact of microproteins 332690, 156126 and 290634 on C. elegans, and featured 3 replicate plates per condition.

One day prior to microprotein stimulation, 60 mm LB NGM + 0.5 mM IPTG agar plates were seeded with 0.5 mL of the relevant bacterial overnight culture (OD600 = 1). Seeded plates were left overnight at room temperature to allow lawns to form and dry and induce microprotein expression. On the day of microprotein stimulation, approximately 150-250 Day 2 adult worms were dispensed onto the seeded plates and incubated at 20°C for 4 hours. Worms were then washed off plates with M9, washed once (RNA-seq 1) or twice (RNA-seq 2) with M9 to remove contaminating bacterial cells. The remaining supernatant was removed, and 200 µL TRIzol (Invitrogen) was added to each cleaned worm pellet. After vortexing the tubes (5 minutes, 4°C), 100 µL HPLC-grade chloroform (J.T.Baker) was added and tubes were shaken by hand (15 seconds). Samples were transferred to Phase Lock Gel Heavy (Quantabio, USA) or standard 1.5 mL (Eppendorf) tubes, and centrifuged (10 minutes, 10,000 x g, 4°C) to separate RNA from denatured proteins, DNA and organic solvents. The RNA-containing aqueous portion was transferred to a new tube, and 1 µL glycoblue co-precipitant (Invitrogen) and 100 µL isopropanol (VWR) were added. Samples were incubated (10 minutes, room temperature), centrifuged (20 minutes, 15,000 x g, 4°C), and the resulting supernatant was removed using an aspirator (Integra Vacuboy). The blue pellet was resuspended in 500 µL ethanol, centrifuged (10 minutes, 15,000 x g, 4°C), and the supernatant was once more removed using an aspirator. The resulting cleaned pellet was resuspended in nuclease-free water and stored at -20°C until use. RNA content was quantified using a Qubit RNA High Sensitivity assay (Invitrogen). Purified RNA was sequenced as summarised in Table 5.

**Table 5.** Summary of RNA sequencing methods.

|  | **RNA-seq 1** | **RNA-seq 2** |
| --- | --- | --- |
| Sequencing provider | Novogene | Francis Crick Institute Genomics STP |
| Library preparation | polyA selection | polyA selection (Watchmaker Genomics) |
| Sequencing platform | Illumina NovaSeq 6000 | Illumina NovaSeq X |
| Output | Paired-end 150 (PE150) | Paired-end 100 (PE100) |

RNA-seq output was processed and analysed using a combination of custom Bash and Python scripts. First, the C. elegans genome and gene annotation files were retrieved from Ensemble (Table 6). STAR ^60^ was used to build, and align reads to, the reference genome. The featureCounts program from the Subread package was used to generate gene expression counts, and DESeq2 ^61^ was used to identify differentially expressed genes (DEG). Gene ontology enrichment was assessed using STRING ^50^.

**Table 6.** Summary of C. elegans reference genome and gene annotation files.

|  | RNA-seq 1 | RNA-seq 2 |
| --- | --- | --- |
| Ensembl release | 112 | 114 |
| Reference genome | Caenorhabditis_elegans.WB<br>cel235.dna.toplevel.fa | Caenorhabditis_elegans.WB<br>cel235.dna.toplevel.fa |
| Gene annotation | Caenorhabditis_elegans.WB<br>cel235.112.gtf | Caenorhabditis_elegans.WB<br>cel235.114.gtf |

## Acknowledgements

The authors would like to thank: Meenakshi Chakraborty and Ann Lin for experimental advice and guidance; Andrew Fire, Ivan Zheludev, Karen Artiles and Dae-Eun Jeong for sharing protocols, materials and advice in support of C. elegans RNA-seq experiments; Aylin Hanyaloglu and Annabelle Milner for experimental advice and guidance regarding host impacts; Kailash Ramlaul and Chris Aylett for the kind gift of the pET9a expression vector; Michael Crone for guidance on Nanopore sequencing; The Genomics STP at the Francis Crick Institute for C. elegans RNA-seq project support.

This research was funded by the Rosetrees Trust (PI DTR: PhD2023\100039 and CF-2023-I-2\112); Rosetrees Trust/Q Charitable Trust (PI DTR: Seedcorn2021\100090), and the Medical Research Council (PI AEXB: MC-A658-5TY30). DTR was funded by a Wellcome Trust/Royal Society Sir Henry Dale Research Fellowship (211230/Z/18/Z).

