## Supplementary Figures 1-8 for "A whole organism screening platform identifies gut microbiome microproteins that modulate host metabolism"

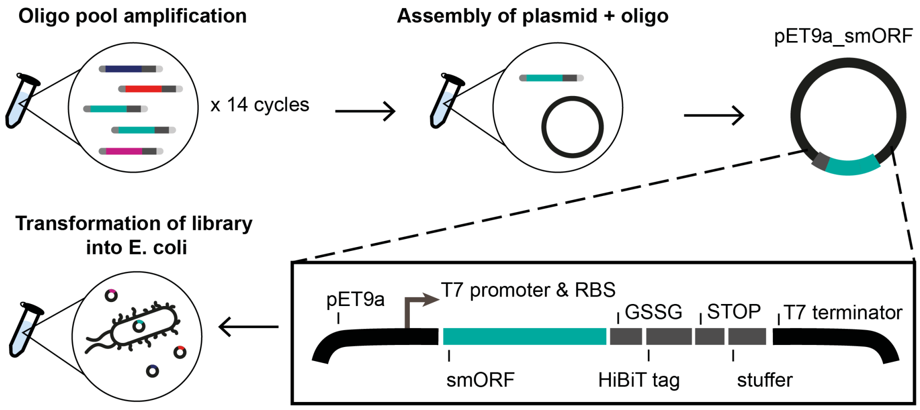


**Supplementary Figure 1. Schematic of cloning strategy**

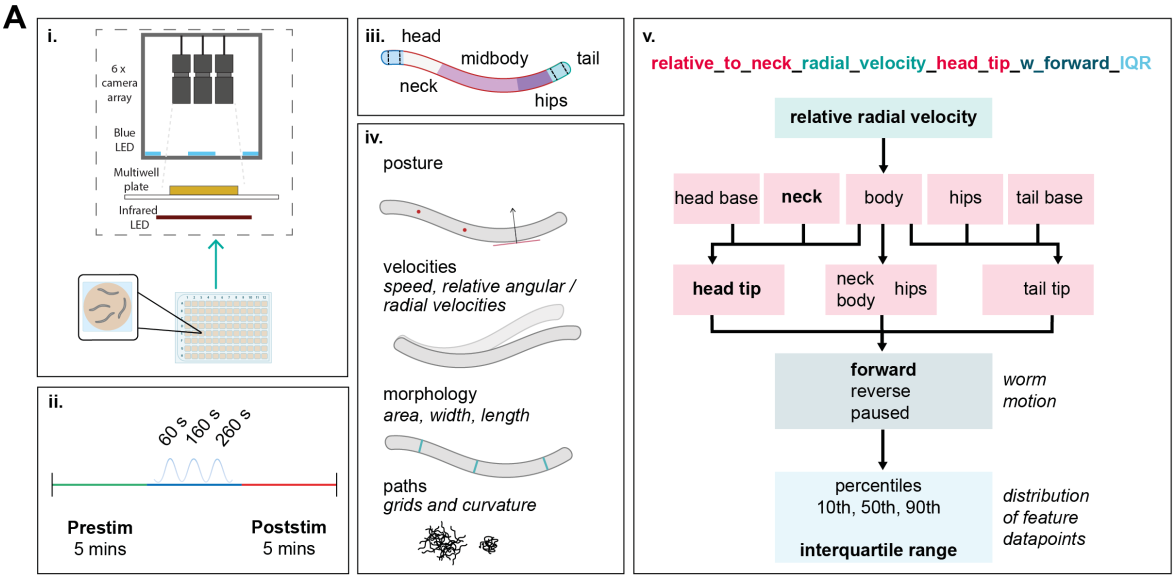


**Supplementary Figure 2. Phenotyping screens in *C. elegans*. i**. Worms in assay microplates are imaged in the Hydra imaging rig. The rig consists of 5 towers, each of which can house one microplate. Each microplate is imaged using six high-resolution cameras, which each image 1/6 of the microplate area. **ii**. A standard imaging run lasts 16 minutes and consists of 3 imaging phases: a prestimulus phase, a bluelight phase with three pulses of blue light, and a post-stimulus phase. **iii**. After image segmentation, each tracked worm skeleton is subdivided into a series of discrete body segments. **iv**. The 3000+ tracked features are all derived from a smaller feature set based on worm posture, velocities, morphology, and path characteristics within and between different body segments. **v**. Break-down on feature naming convention.

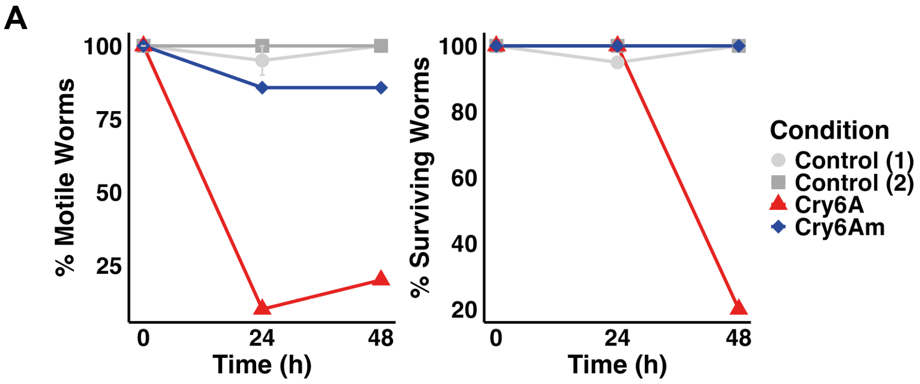


**Supplementary Figure 3. Screening of Cry6A and Cry6Am impacts on worms.**

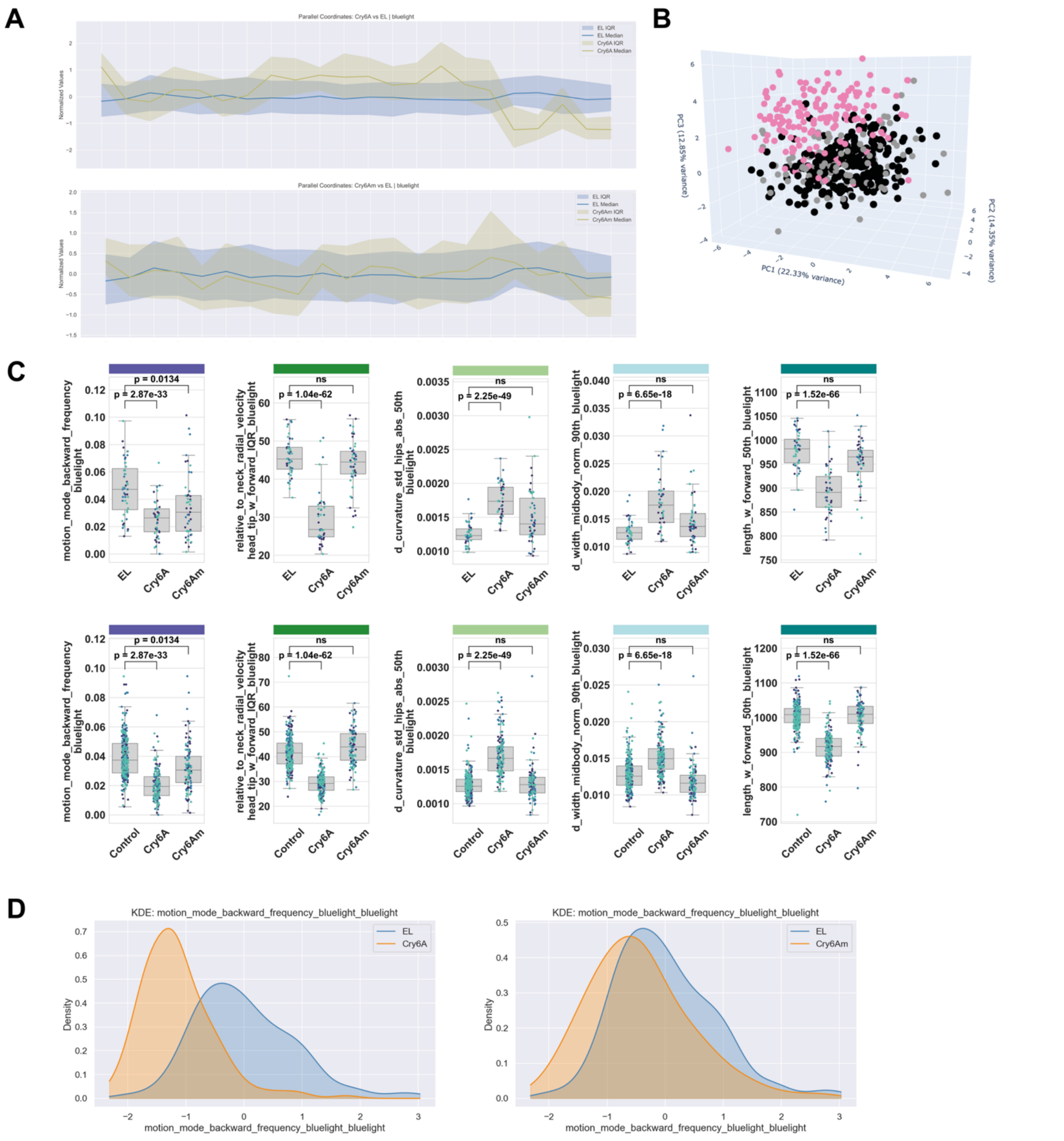


**Supplementary Figure 4. Impact of Cry6A and Cry6Am expression on *C. elegans* behaviour in Screens 2 and 3. A**. Parallel coordinate plot showing normalized scores across Tierpsy 22 feature set for i. Cry6A vs. *E. coli* control and ii. Cry6Am vs. *E. coli* control in Screen 3. The median score per feature is represented by the line, and the interquartile range is represented by the shaded area. **B**. Principal component analysis plot of feature scores from Screen 3. Each point represents a unique well containing an average of 3 worms. Conditions shown are Cry6A (pink), Cry6Am (grey) and *E. coli* control (black). **C.** Boxplots illustrating the distribution of feature scores from select features from Screen 2 (first row) and Screen 3 (second row). Each point represents the average feature score across all worms in a single well, and is coloured according to the screening day it is derived from. For each feature, scores from the control group were compared to those from each environmental condition using univariate two-sample t-tests. P-values were corrected for multiple comparisons using the Benjamini–Yekutieli procedure to control the false discovery rate. ns: not significant ($p > 0.05$). **D**. Kernel density estimation (KDE) plot of representative motion feature from Screen 3, showcasing difference in feature distribution between Cry6A, Cry6Am and *E. coli* Lemo21 controls. Number of wells screened in Screen 2 and 3: Cry6A: 43 and 159; Cry6Am: 49 and 104; Control: 47 and 264.

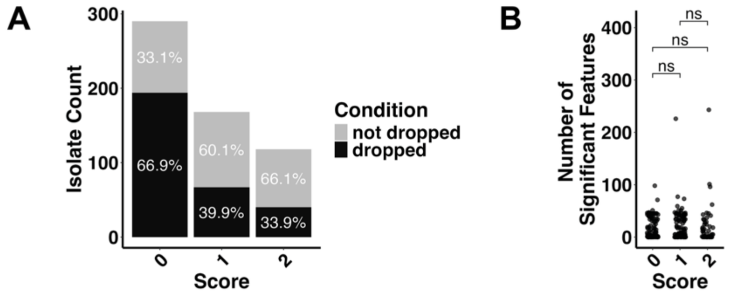


**Supplementary Figure 5. Correlation between lawn growth score and tracking output.** Each isolate from the arrayed smORF library was grown on LB agar with 0.5 mM IPTG overnight at room temperature, and the resulting lawn was assigned a score depending on the quality of the lawn (0: significant growth impact, absent or minimal lawn; 1: intermediate growth impact, thin lawn, 2: minimal growth impact, thick lawn). **A**. The number isolates that were dropped/not dropped from Screen 1 analysis, grouped by growth score. The percentage of isolates that were dropped/not dropped per growth score is shown. A Spearman's rank correlation analysis confirmed that there was a significant moderate negative correlation between the growth score of a well code and whether it was dropped from the analysis (Spearman Correlation Coefficient: -0.306, p-value: $5.574 \times 10^{-14}$ . **B**. The number of significant features impacted by a given isolate from Screen 1, and its corresponding growth score. Each point represents a single well code (i.e. isolate) from the arrayed library. A Kruskal-Wallis test was used to determine the statistical significance of differences between the groups. ns: not significant ($p > 0.05$).

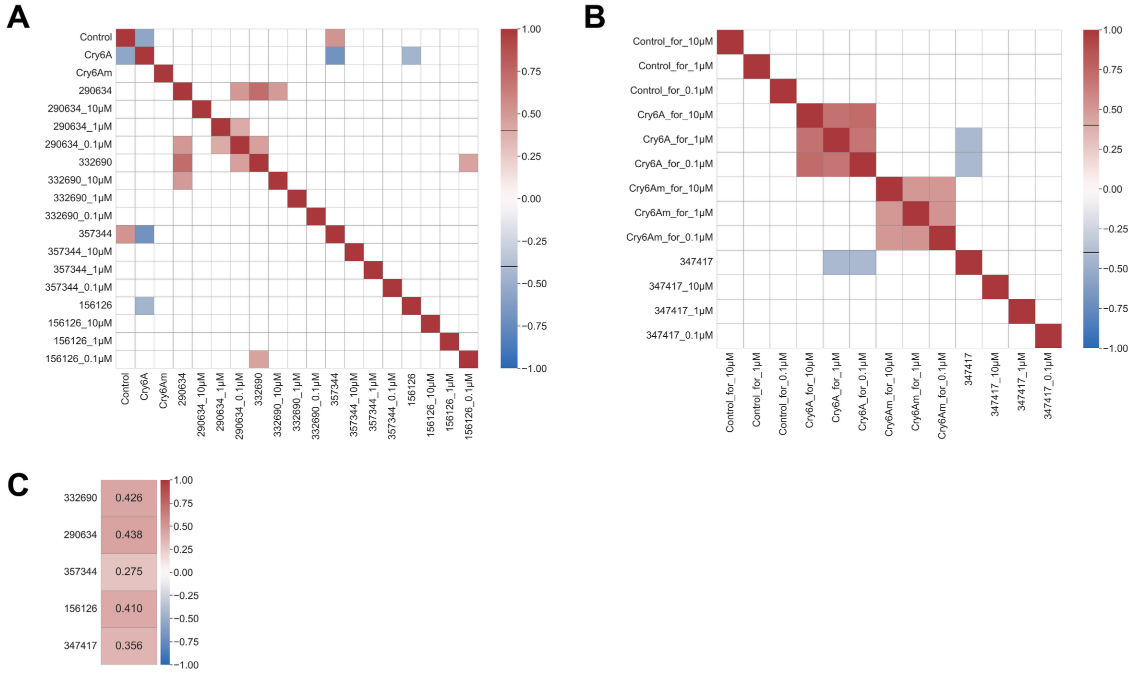


**Supplementary Figure 6. Correlation between impact of recombinant and synthetic microprotein pairs on *C. elegans* behaviour. A-B**. Spearman rank correlation matrices between samples based on Z-normalised mean feature values. Correlation coefficients with absolute values < 0.4 (as indicated in the figure colour scale) are not displayed. Microprotein IDs correspond to recombinant microproteins, whilst IDs followed by a concentration correspond to the concentration of synthetic microprotein. **A**. Samples and controls associated with synthetic microproteins resuspended in water. B. Samples and controls associated with synthetic microprotein resuspended in isopropanol. C. Spearman correlation coefficients describing the relationship between the phenotypic impact of recombinantly expressed microproteins and the corresponding synthetic microproteins, where synthetic profiles represent the mean across the three tested concentrations. Correlations were calculated using Z-normalised mean feature values, as in A-B.

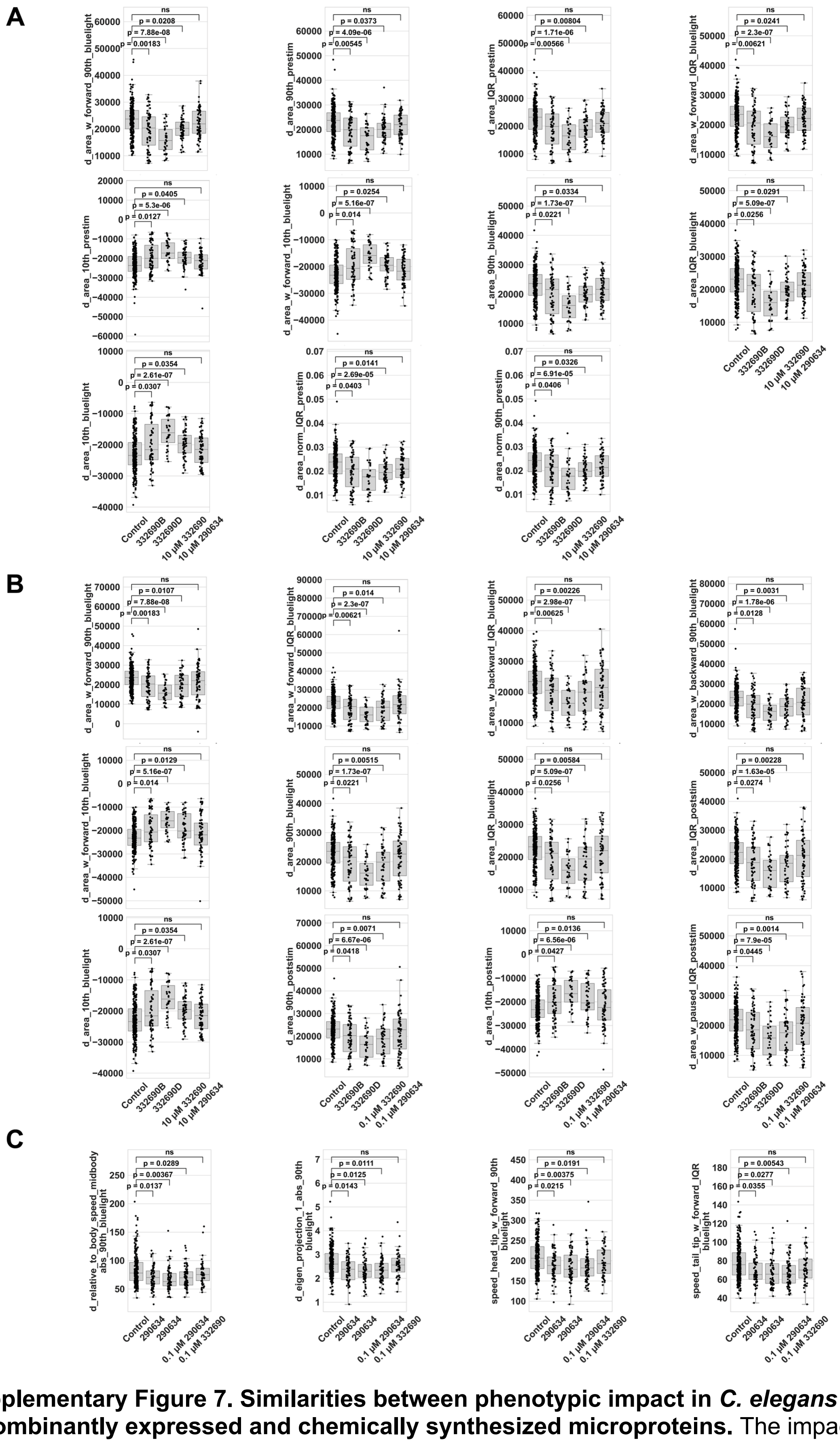

**Supplementary Figure 7. Similarities between phenotypic impact in *C. elegans* of recombinantly expressed and chemically synthesized microproteins.** The impact of recombinantly expressed microprotein 332690 and 290634 was compared against that of 10 µM, 1 µM and 0.1 µM of their chemically synthesized counterparts. A standard behavioural screen was carried out, with the exception that worms were fed on a lawn of control strain *E. coli* to which a solution of synthetic microprotein in solution had been added. After 4 hours of exposure, worm behaviour was recorded and analysed using Tierpsy tracker and custom Python analysis scripts. Boxplots comparing impact of recombinant and synthetic Microprotein **A.** 332690 and **B**. 290634. All features that were significantly impacted by both isolates of recombinantly expressed microprotein and at least one concentration of synthetic protein are shown. Each point corresponds to the average feature score across a single well, containing approximately 5-10 worms. For each feature, scores from the control group were compared to those from each environmental condition using univariate two-sample t-tests. P-values were corrected for multiple comparisons using the Benjamini–Yekutieli procedure to control the false discovery rate. ns: not significant ($p > 0.05$). Number of replicate wells per condition: Control: 264; 332690B: 66; 32690D: 32; 10μM_332690: 55; 0.1μM_332690; 10μM_290634: 67; 0.1μM_290634: 69.

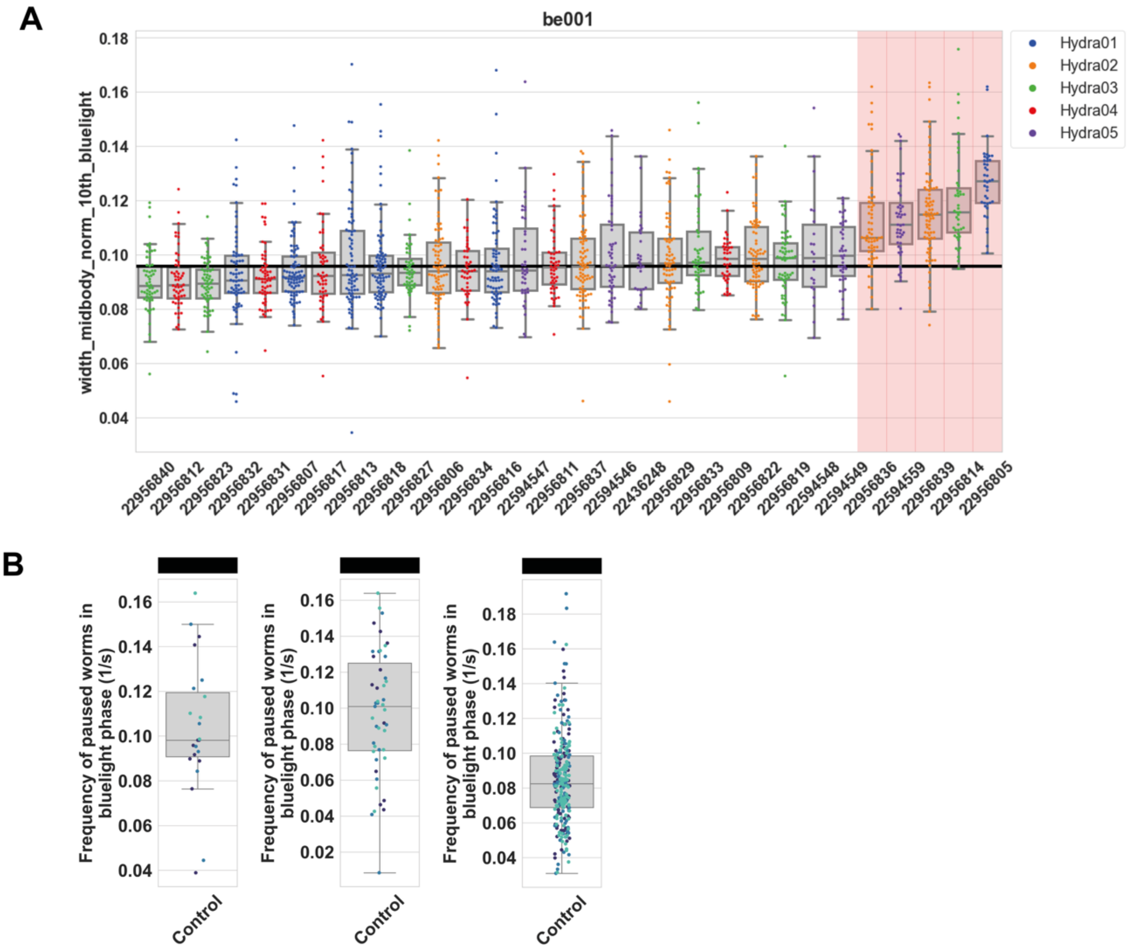


**Supplementary Figure 8. Phenotyping screen quality control.** **A.** Data from recordings deemed to be out of focus are removed from phenotyping screen analyses. For a given width feature, all data from cameras whose median score differs by 2 standard deviations or more from the overall median score (represented by the shaded right rectangle) are discarded. The boxplots illustrate the distribution of feature scores, where each well corresponds to the average feature across a single well (containing an average of 3 worms) and are coloured according to the camera that recorded that well. **B**. To rule out any bias in screen results as a result of variation between the 3 days of screening, we examine the distribution of feature scores across all screening days for a given feature-set for three successive screens (i, ii, iii). The boxplots illustrate the distribution of feature scores for a representative motion feature. Points correspond to the average feature score across a single well (containing an average of 3 worms), and are coloured according to the screening day. Number of replicate wells screened: i. 23, ii. 47, iii. 264.
